# Characterization of DNA processing protein A (DprA) of the radiation-resistant bacterium *Deinococcus radiodurans*

**DOI:** 10.1101/2022.07.25.501491

**Authors:** Dhirendra Kumar Sharma, Hari S. Misra, Yogendra S. Rajpurohit

## Abstract

The uptake of environmental DNA (eDNA) by certain bacteria and its integration into their genome creates genetic diversity and new phenotypes. A DNA processing protein A (DprA) is part of a multiprotein complex and facilitate natural transformation (NT) phenotype in most bacteria. The *Deinococcus radiodurans,* an extremely radioresistant bacterium, is efficient in N T, and its genome encodes nearly all the components of the natural competence complex. Here, we have characterized the DprA of this bacterium (DrDprA) for the known characteristics of DprA proteins in other bacteria and the mechanisms underlying its roles in the transformation of eDNA into the bacterial genome. DrDprA is found to be a unique domain organization implicating some unique functions compared with DprA of other bacteria. *In vitro* studies showed that the purified recombinant DrDprA binds to both ssDNA and dsDNA with nearly equal affinity and protects ssDNA from nucleolytic degradation. DrDprA showed a strong interaction with DrRecA indicating its role in RecA catalyzed functions *in vivo*. Mutational studies identified amino acid residues responsible for its oligomerization, interaction with DrRecA, and DNA binding characteristics of DrDprA. Further, we demonstrated that both oligomerization and DNA binding properties of DrDprA are integral to its support in DrRecA catalyzed strand exchange reaction (SER) *in vitro.* These results suggested that DrDprA is largely structurally conserved with DprA homologs but showed some unique structure-function features like additional domain, the same affinity to ss/ds DNA and both oligomerization and DNA binding domains collectively contribute to its support in DrRecA functions.

## Introduction

Natural competence is a genetically regulated mode of horizontal gene transfer to acquire the extracellular DNA through transformation. Several bacteria show natural transformation (NT), through which they acquire external DNA from the environment and recombine it with their genetic material to introduce genetic diversity [1–5] and fitness of any bacterium under adverse environmental conditions through different modes including using DNA as foodstuffs [6, 7] and utilization of the extracellular DNA to facilitate recombination repair of damaged DNA [1, 5, 8]. Mechanistically, it involves two major steps; (1) uptake of external DNA into the cytosol by a macromolecular complex and (2) the integration of transforming DNA into the host chromosome by homologous recombination (if transforming DNA is chromosomal DNA) or stabilization of transforming DNA into a functional circular plasmid DNA by single-strand annealing (SSA) activity (if transforming DNA is plasmid DNA) [9]. The DNA uptake system comprises either type II secretion systems or type IV pili (T4P) [6, 10, 11]. The components of the uptake complex have been identified, and their roles in DNA uptake have been characterized. For instance, the external dsDNA translocated into the cytoplasm is first converted into ssDNA by EndA nuclease and then processed by the combined action of receptor protein ComEA (permease), transmembrane channel protein ComEC, and the ATPase motor protein ComF [7, 12–14]. The internalized ssDNA is protected by single-stranded DNA binding protein (SSB) and by a natural transformation-specific recombination mediator protein (DprA) [15–17]. The DprA protein further facilitates the RecA recombinase loading on incoming ssDNA by overcoming the SSB protein barrier [16, 18–20]. Finally, transforming DNA is integrated into host genetic materials through homologous recombination [6, 9, 21]. DprA has been found in almost all bacteria, including *Deinococcus radiodurans* but is functionally characterized in a few. It is also named CilB or Smf in different bacteria. DprA is also required to load RecA on SSB-coated ssDNA in different bacteria [16–20].

*D. radiodurans* is best characterized for its extraordinary resistance to DNA damaging agents, including radiation and desiccation [22, 58]. The cytogenetic features of this bacterium are unique with its ploidy in a multipartite genome system, and the existence of tetrad colonized cell phenotypes [22, 23, 58]. A possible translocation of an undamaged copy of genome elements or its constituents from one cell of the tetrad to other cells and pumping out of damaged materials from the cells has been hypothesized for the efficient DSB repair and radioresistance in this bacterium [23, 24]. The molecular basis of transporting such materials across the cells and from the outside environment is not well understood. However, this bacterium exhibits NT and has acquired nearly 10% of its genetic materials from other organisms and thus has created genetic diversity [25, 26]. Therefore, the possibility of NT contributing to its extreme phenotypes, albeit indirectly, cannot be ruled out and would be worth investigating. The *D. radiodurans* genome encodes the components of the NT system, including DprA (DrDprA). It has been shown that the RecFOR complex / DdrB protein partially compensates for the functional requirement of DrDprA in the DNA transformation process [27]. The molecular basis of DrDprA that makes it different from the DprA of other bacteria would require detailed studies on its structure and functions. Here, we report the structure-function characterization of DrDprA, a core component of the NT system. We demonstrated that DrDprA exhibits an almost equal preference for binding to both ssDNA and dsDNA. The conserved residues for the oligomerization and RecA interaction have been identified and confirmed through mutagenesis. DrDprA could support the RecA functions and reduce the SSB interference in the DNA strand exchange activity of DrRecA. It was able to protect DNA from nuclease degradation, and its deletion mutant was found to be defective in at least plasmid transformation. Taken together, these results suggested that DrDprA has all the DNA metabolic properties that are known in other bacterial proteins and are required for the transformation of external DNA in this bacterium.

## Results

### 1) DrDprA of *D. radiodurans* has conserved the structural domains of DprAs in other bacteria

DprA protein size from different bacteria varies from 240 *(Campylobacter jejuni)* to 398 amino acids (AA) *(Synechocystis* and *Neisseria gonorrhoeae)* [37]. The DprA protein of *D. radiodurans* (DrDprA) is 370 amino acids long [25]. DprA proteins from different organisms showed significant similarities at the amino acid sequence levels. The multiple sequence alignment (MSA) of DrDprA with DprA of *Rhodopseudomonas palustris, Helicobacter pylori,* and *Streptococcus pneumoniae* showed 31 to 38 % identity (Fig. 1). Structural studies of DprA from *R. palustris* (PDB-3MAJ), *S. pneumoniae* (PDB-3UQZ) [19] and *H. pylori* (PDB-4LJK) [38] have revealed that protein is structurally divided into a central <u>R</u>ossmann <u>F</u>old (RF) domain, N-terminal <u>S</u>terile <u>A</u>lpha <u>M</u>otif (SAM) and C-terminal *<u>D</u>rosophila melanogaster* <u>M</u>iasto <u>L</u>ike protein <u>1</u>(DML1) domain. MSA showed that DrDprA also contains all three domains; the SAM domain ranges from 3-78 amino acids, the RF domain from 80-296 amino acids, and the DML1 domain from 308-367 amino acids (Fig. 1). However, MSA revealed diversity in domain organization amongst DprA homologs. For instance, the SAM domain is generally associated with the RF domain in most DprA except *H. pylori* and *Pyrococcus furiosus,* while the DML1 domain is found in the DprAs of *Mycobacterium tuberculosis, D. radiodurans, Vibrio cholera, R. palustris, Neisseria meningitides, Haemophilus influenzae,* and *Synechocystis sp.* (Fig. 1A). It has been shown that RF domain contributes in protein-protein interaction and higher-order oligomerization while N-terminal SAM and C-terminal DML1 domain may contribute in DprA functions by assisting oligomerization and protein-protein interaction [18, 19]. In eukaryotes, SAM domain-containing proteins have been shown to regulate several developmental changes [39] while the DML1 domain is considered Z-DNA binding domain [40].

**Figure 1.**
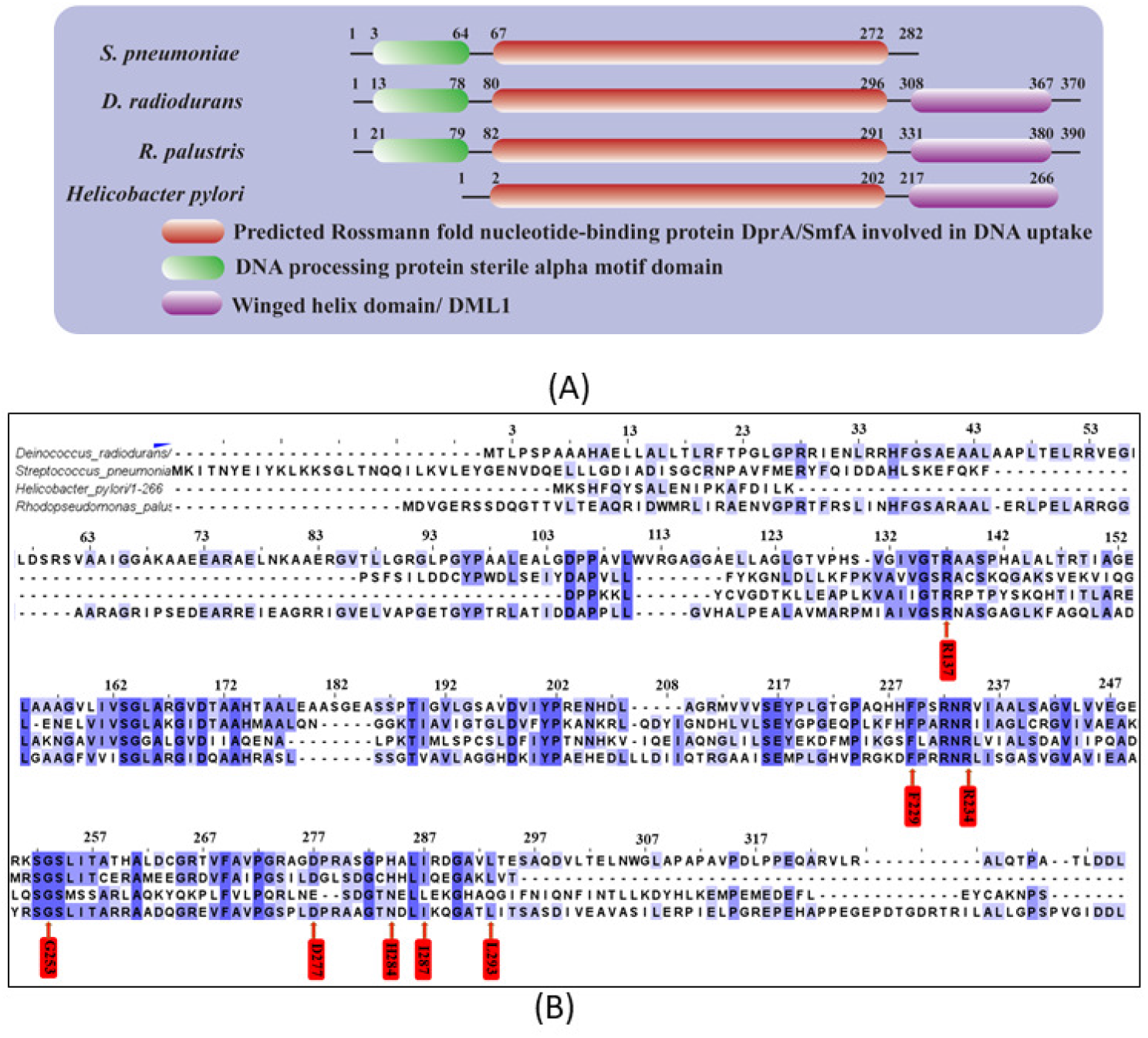
Domain architecture of DprA. (A) Domain architecture of DprA protein of selected bacteria (*S. pneumoniae*, *R. palustris, D. radiodurans* and *H. pylori*). Central Rossmann fold (RF) is conserved and present in all DprA proteins while N-terminal SAM domain common in all DprA except *H. pylori*. An additional C-terminal DML1 domain present in DprA of *D. radiodurans, and R. palustris.* (B) Multiple Sequence alignment of DprA peoteins. Red arrow represent conserved putative amino acids responsible for the DprA-RecA interaction (I287, G253, and D277), DprA-DprA interaction (L293, and H284), and interaction with DNA (R137, R234, and F229) as predicted *in silico* analysis.

### 2) DrDprA forms a nucleoprotein complex with ssDNA and dsDNA

The recombinant DrDprA protein was purified to near-homogeneity as described in methods (Fig. S1, for reviewers’ information only) and its DNA binding activity was monitored for both dsDNA and ssDNA. Since the DNA binding ability of DprA homologs has been shown to increase with the increasing length of DNA upto 80 bp [20, 41], the DNA length requirement of the DrDprA for its binding was evaluated with different size substrates. It was found that DprA forms an unstable NPC (nucleoprotein complex) with DNA smaller than 40 bp and a stable NPC with 167bp ssDNA /dsDNA (Fig. 2A). Interestingly, DrDprA could bind with both ssDNA and dsDNA (167bp) with equal binding affinity. The Kd values for ssDNA was found to be 3.039 ± 0.09724μM and 3.526 ± 0.1172 μM for dsDNA (Fig. 2B, D and E). Competition experiments further ascertained the binding preference of DrDprA with ss/ds DNA substrates. Results showed that the DrDprA has the same affinity for binding to ssDNA and dsDNA as the NPC of both radiolabeled ssDNA and dsDNA could not be destabilized by 50 to 400 molar excess of cold ssDNA and/or dsDNA and vice versa (Fig. 3). This characteristic of DrDprA is found to be unique when compared with DprAs of other bacteria. For example, the DprA proteins of *S. pneumoniae* (SpDprA) and *Bacillus subtillis* (BsDprA) show binding with ssDNA only [17, 19, 20]. At the same time, the DprA of *H. pylori* (HpDprA) interacts with both ssDNA and dsDNA; it modestly prefers ssDNA over dsDNA [42]. DrDprA also showed interaction with circular plasmid DNA (M13mp18) incubated with and without ATP, indicating that DrDprA does not require DNA ends to bind with DNA (Fig. 2C, F). A similar observation on DprA binding to DNA independent of open ends has been reported for the SpDprA and HpDprA [20, 42].

**Figure 2.**
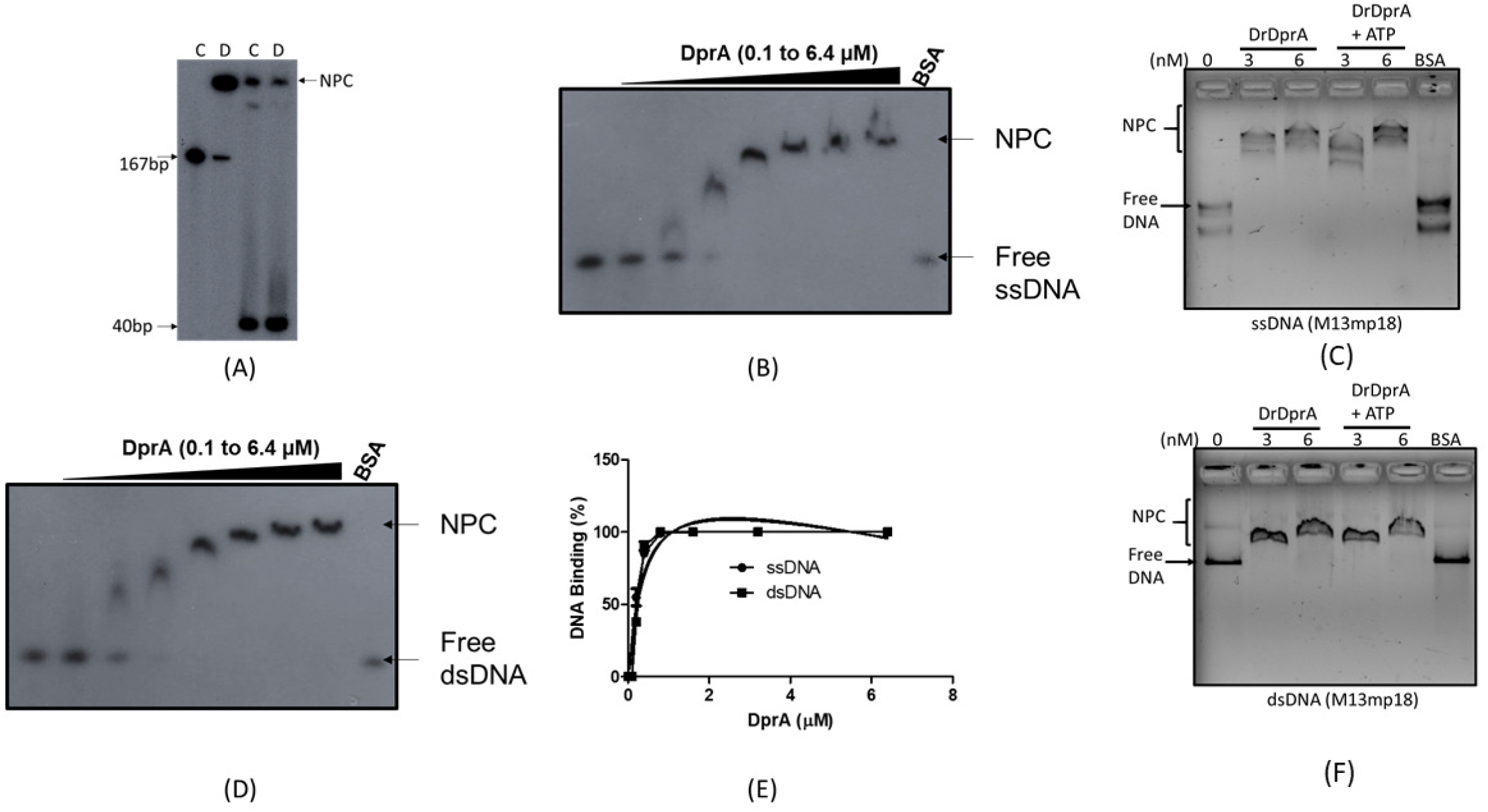
DNA binding activity of DrDprA. Recombinant DrDprA proteins was purified and incubated with [^32^P] labeled (A) single strand DNA of different size (40mer and 167mer), (B, D) 167mer ssDNA /dsDNA with increasing DrDprA concentration (0.1, 0.2, 0.4, 0.8, 1.6, 3.2, and 6.4μM) in a reaction buffer as described in methods. Reaction mixtures were separated on 8% native PAGE, gels were dried and autoradiogram done. (E) bound DNA band intensities were quantified densitometrically and percent bound fractions were calculated. Results were plotted and analyzed using Graphpad Prism software. (C, F) DrDprA binding with M13mp18 ss and dsDNA monitored on agarose gel (0.8%). Data shown are representatives of the reproducible experiments repeated two times.

**Figure 3.**
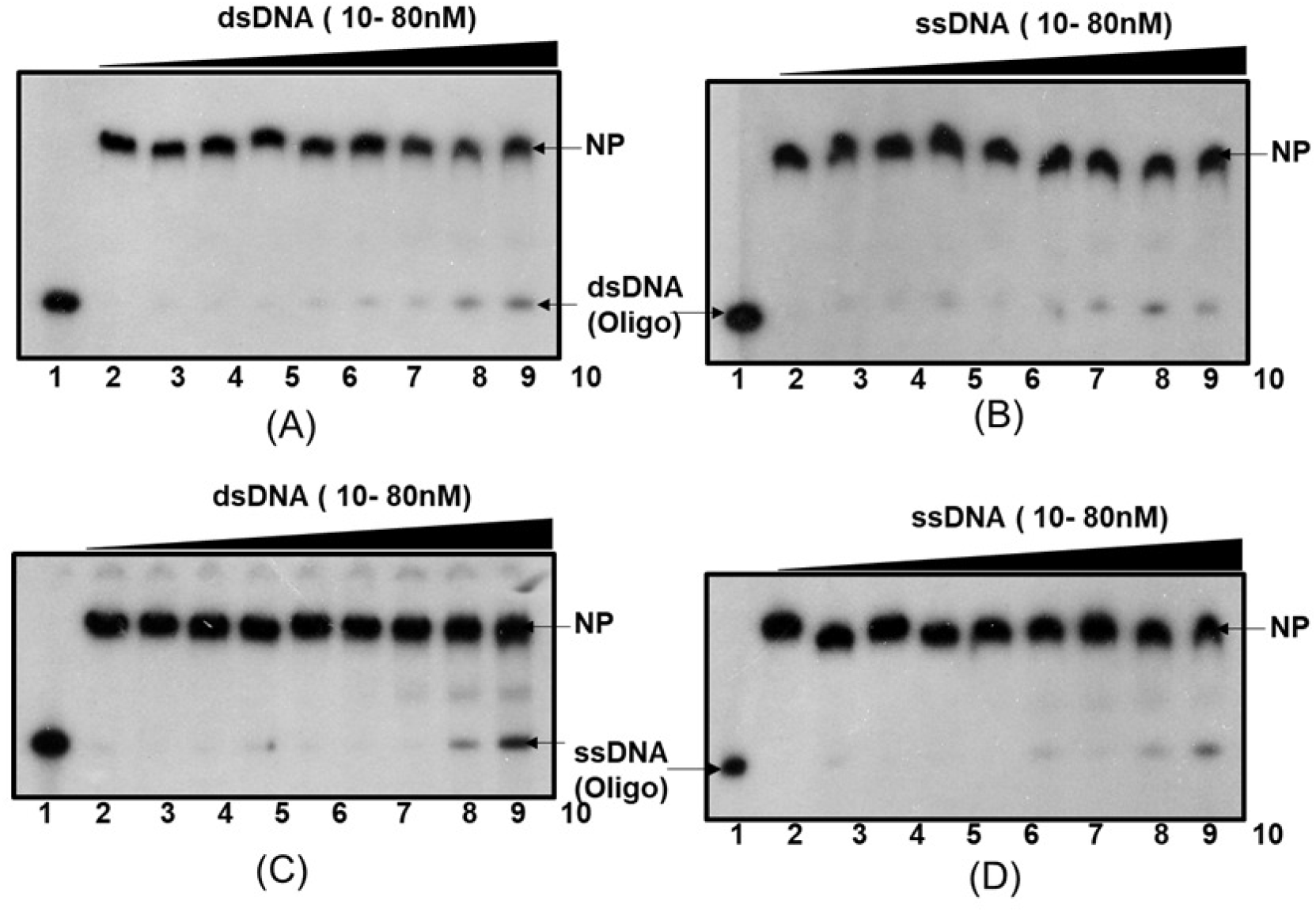
DrDprA DNA binding preference between ssDNA and dsDNA substrates. 1.6 μM of purified recombinant DrDprA protein incubated with [^32^P]-labeled dsDNA (**A, B**) and ssDNA (**C, D**). Protein bound to these substrates were chased with increasing concentration of unlabeled ssDNA (**B, D**) and dsDNA (**A, C**). Mixtures were analyzed on native PAGE and autoradiograms from a reproducible representative experiment is shown. Both free DNA and DNA bound to protein were quantified densitometrically from three independent experiments and the percent bound fractions were calculated.

The binding of deinococcal proteins that prefers dsDNA otherwise to ssDNA when compared with their homologs in other bacteria has been observed earlier. A notable example is the preference of dsDNA by deinococcal RecA catalyzing inverse strand exchange reaction as shown earlier [33, 43, 44]. These results suggested that DrDprA can bind with both ssDNA and dsDNA in a sequence and DNA open ends independently. Further, the ability of DrDprA to bind with dsDNA is an interesting phenotype and suggests a possible role of this protein in DNA metabolism beyond natural transformation. The salt and temperature stability of DrDprA NPC with ssDNA and dsDNA was evaluated, and it found that up to 250mM NaCl concentration, more than 90% NPC was retained (data not given). The temperature stability of DrDprA or its nucleoprotein complex (NPC) with ssDNA was estimated at up to 45°C. Beyond this temperature, the NPC dissociate rapidly irrespective of DNA binding order and incubation at different temperature (Fig. S2, for reviewers’ information only).

### 3) DrDrpA interacts with DrRecA

The role of some DprAs in the loading of RecA to ssDNA has been reported. Further, the interaction of SpDprA with SpRecA has also been shown [17, 18, 20, 45, 46]. Therefore, the interaction of DrDprA with the DrRecA protein was monitored *in vitro, ex vivo,* and *in vivo.* For *in vitro,* the DrRecA and DrDprA interaction was monitored with purified proteins by Surface Plasmon Resonance (SPR) as described in the methods. The results showed a concentrationdependent increase in the SPR signal when purified DrDprA was brought in contact with DrRecA in solution (Fig. 4C). A strong interaction was supported with a suitable Lorentz fit and Kd value of 2.93 × 10^-7^ ± 1.34 × 10^-8^ Molar for DrDprA interaction with DrRecA. The *ex vivo* interaction of these proteins was checked in surrogate *E. coli* BTH101 using a bacterial two-hybrid system. For that, the T18 tagged DrDprA and T25 tagged DrRecA were co-expressed in *E. coli* BTH101 (Fig. 4A, B). The reconstitution of active adenylate cyclase (CyaA) by joining the T18 and T25 domains of CyaA, upon the interaction of these two proteins, was monitored as the expression of the β-galactosidase enzyme [39]. As expected, the cells co-expressing T18-DrDprA and T25-DrRecA produced intense blue-colored colonies on the X-gal plate, which was not observed in cell co-harboring T18 and T25 expressing vectors as negative control (Fig. 4A). The level of expression of β-galactosidase activity was very high as predicted from the blue color intensity of the bacterial colonies and activity measured in solution suggesting a strong interaction between DrRecA and DrDprA *ex vivo* (Fig. 4A). The interaction of DrDprA and DrRecA was also monitored *in vivo*. For that, DrRecA tagged with (his)^6^, and DrDprA tagged with T18 domain of CyaA were expressed in *D. radiodurans.* Prospective interaction was monitored by coimmunoprecipitation (Co-IP) using histidine antibody, and the partner was detected with T18 antibody. The results showed that lanes representing samples from either DrDprA or DrRecA did not show a signal with the T18 antibody (Fig. 4B). At the same time, those from cells coexpressing his-DrRecA and T18-DrDprA produced a band of molecular weight of ~56 kDa which is theoretically the size of T18-DrDprA (Fig. 4B). Together, these results suggested that DrRecA with DrDprA proteins interact physically in *D. radiodurans*.

**Figure 4.**
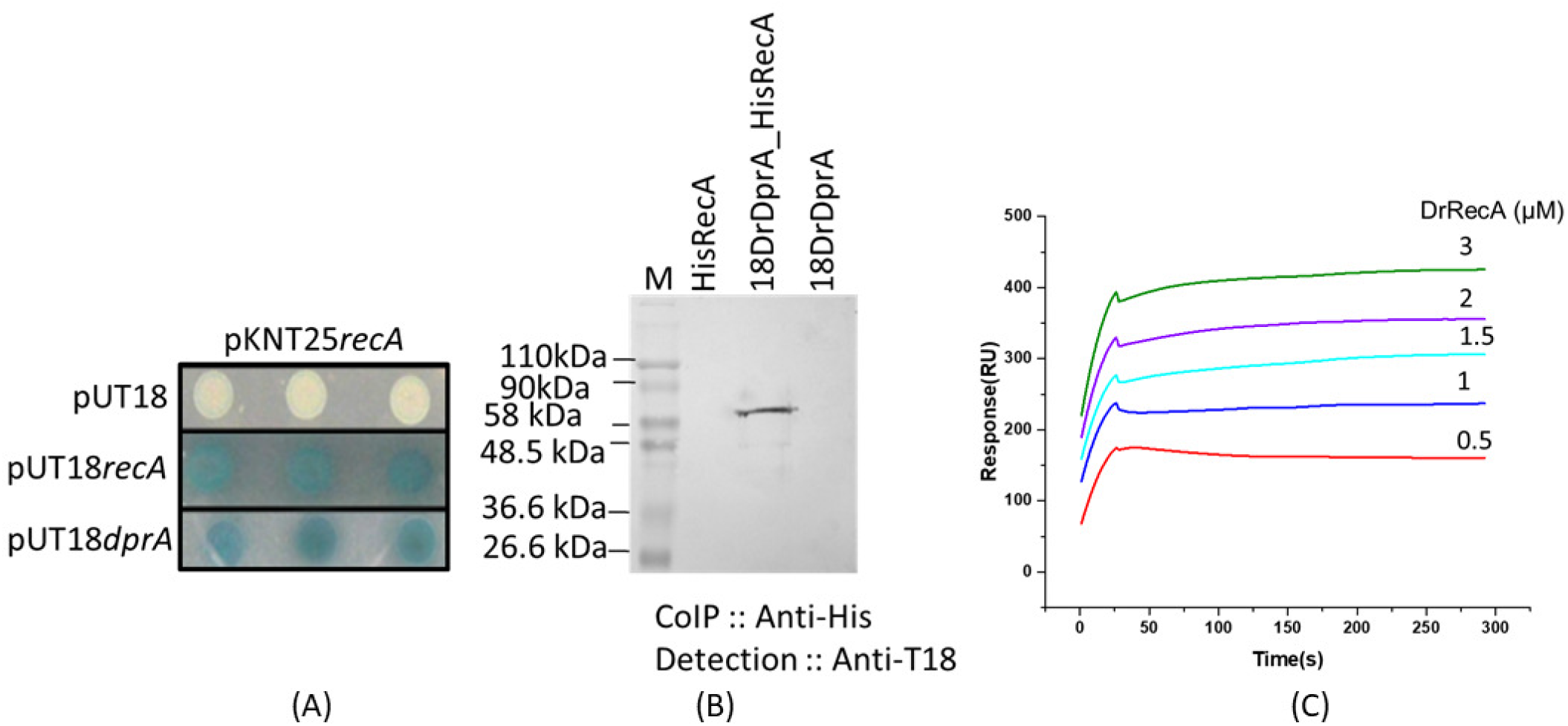
DrRecA and DrDprA interaction in a surrogate *E. coli (ex vivo),* in *D. radiodurans* cells (*in vivo*), and *in vitro.* (A) T18 and T25 tags of adenylate cyclase alone and tagged with *dprA* and *recA* gene of *D. radiodurans* were cloned in BACTH plasmids. These plasmids were transformed into an *E. coli* BTH101 host. The interaction of proteins tagged with T18 and T25 were monitored as white-blue colonies. RecA-C18 and RecA-C25 were used as positive control while C18 and C25 tags expressing cells were used as a negative control. (B) Cell-free extracts of *D. radiodurans* cells co-expressing C18-DrDprA and His-DrRecA from pVHSM and pRAD plasmid used for immunoprecipitation assay respectively. An immunoprecipitation done using anti-His antibody and immunoprecipitates were separated on SDS-PAGE followed by immunoblot detection using antibodies against T18 domain of CyaA (Immunoblot Anti-T18) as detailed in Materials and Methods. (C) 2.5μM DrDprA protein was immobilized on gold sensor chip followed by incubation with different concentration (0.5–3 μM) of the DrDrecA (**C**). Surface Plasmon Resonance (SPR) signals were recorded and data was processed as described in methods. Data given are representative data from the experiments repeated three times.

### 4) DNA binding and oligomerization ability of DrDprA is required for its support to RecA functions

DprAs of other bacteria is known to improve the DNA strand exchange reaction (SER) of RecA [18, 20, 46]. DrDprA showed strong physical interaction with DrRecA (Fig. 4). Therefore, the support of DrDprA in the DNA strand exchange activity of DrRecA was evaluated using an oligo-based (short homology) and M13mp18 DNA-based (extended homology) DNA strand exchange reactions. The low efficiency in SER was observed at 0.5μM DrRecA protein in oligo-based reaction, which had significantly improved upon addition of 0.5 to 4 μM DrDprA (Fig. 5A). Similarly, the addition of 2μM of DrDprA has substantially improved the M13mp18 DNA-based SER by DrRecA (Fig. 5B). The improvement of SER in the presence of DrDprA is in agreement with the similar effect of DprA in recombination reaction in other bacteria. These results suggest that the presence of DrDprA has positively impacted the SER by DrRecA.

**Figure 5.**
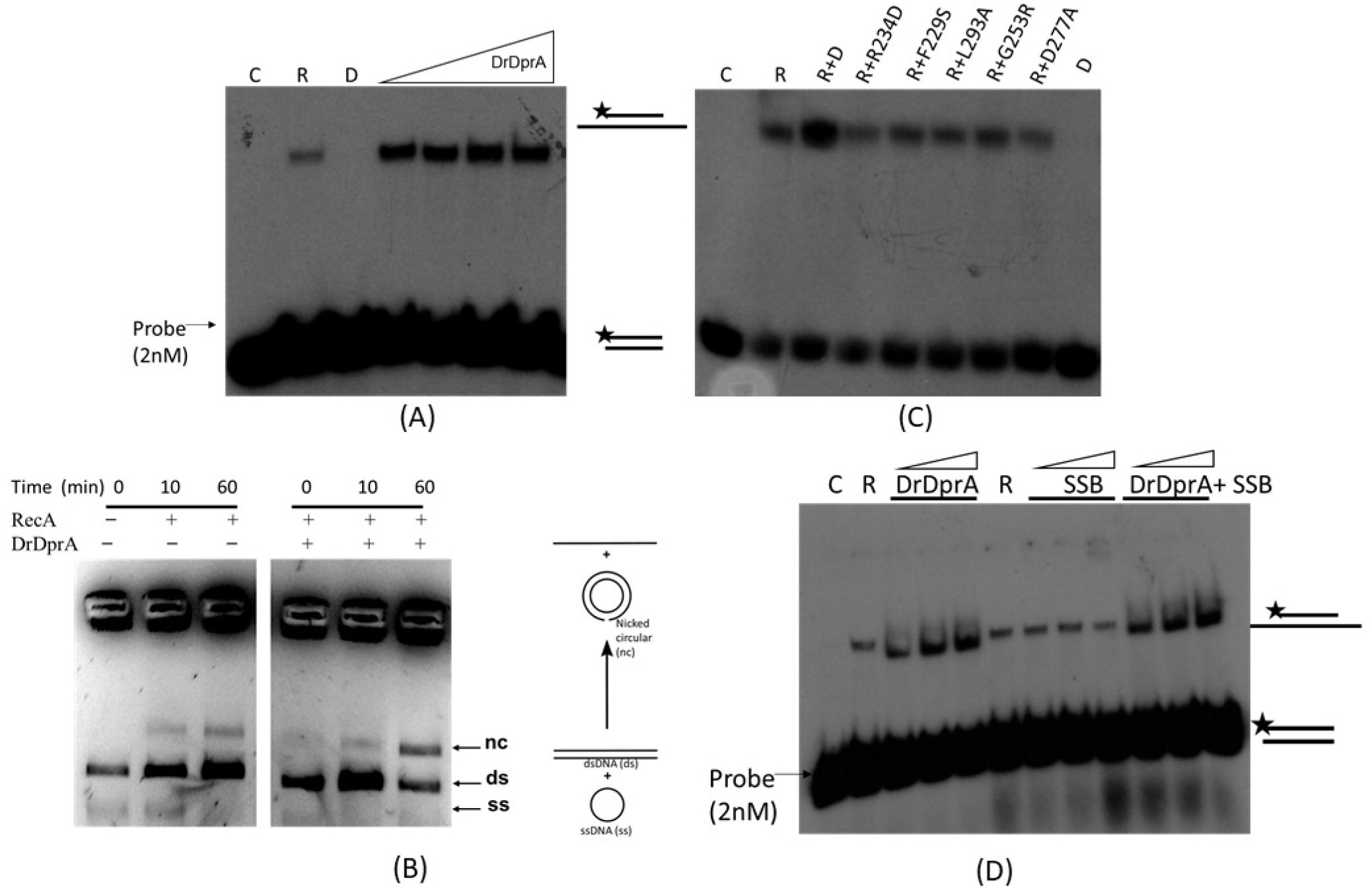
Stimulatory effect of DrDprA on the DrRecA catalized DNA strand exchange activity and DrDprA role in limiting SSB inhibitory effect. (A) In oligo-based DNA strand exchange activity, 0.5μM of DrRecA protein was used. Increasing concentration of DrDprA (0.5, 1, 2, and 4μM) was used to see stimulatory effect of DrDprA. (B) M13mp18 based DNA strand exchange reaction carried out as detailed in methods. DrRecA and DrDprA concentration used are 2.5μM and 2μM, respectively. (C) wild type DrDprA and its different mutant proteins were tested for their stimulatory effect on DrRecA catalized oligo-based DNA strand exchange activity. (D) DrDprA role in limiting the SSB inhibitory effect during DNA strand exchange. DrDprA concentration used are 1, 2, and 4μM while SSB concentration used were 0.3, 0.6, and 1.2μM. C-buffer control reaction, R-DrRecA, and D-DrDprA. Results shown is a representative data of the reproducible experiments repeated three times.

DrDprA is a DNA binding protein that undergoes oligomerization *in solution* and interacts with DrRecA. So, the involvement of these attributes in the stimulation of SER of DrRecA was studied. Earlier, the key amino acids involved in the protein-protein interaction between DprA and RecA have been identified [19, 20, 47]. Based on these reports, the amino acid sequence of DrDprA was mapped with DprA of *S. pneumonia, R. palustris,* and *H. pylori,* and corresponding amino-acids residues of DrDprA were mapped. The putative amino acids responsible for the DprA-RecA interaction (I287, G253, and D277), DprA-DprA interaction (L293, and H284), and interaction with DNA (R137, R234, and F229) were predicted *in silico* (Fig. 1B). The site-directed mutants of these sites were generated, the recombinant proteins were purified (Fig. S1, E, for reviewers’ information only) and checked for their corresponding functions *in vitro.* As predicted, the R137, R234, and F229 mutant lost DNA binding activity with a 167bp long substrate and also showed altered oligomerization profile on size exclusion chromatography compare to wild type (Fig. 6 and S3, for reviewers’ information only). The predicted oligomer mutants (L293 and H284) showed reduced DNA interaction compared to wild-type DrDprA and defect in olizomerization (Fig. 6 and S3, for reviewers’ information only). However, DprA-RecA interaction mutants (G253 and D277) are proficient in DNA interaction similar to wild-type DrDprA and maintain wild type DrDprA profile of olizomerization (Fig. 6 and S3, for reviewers’ information only). When these mutants were tested for their support in SER function of DrRecA, the stimulatory effect of G253R and D277A (RecA-DprA interaction) mutants, L293A (oligomer mutant) and R234D and F229S (DNA binding) mutants on SER of DrRecA was completely abolished (Fig. 5C). This suggested that in addition to DrDprA-DrRecA interaction, the oligomerization and DNA binding properties of DrDprA are also crucial for its recombination mediator protein function with DrRecA.

**Figure 6.**
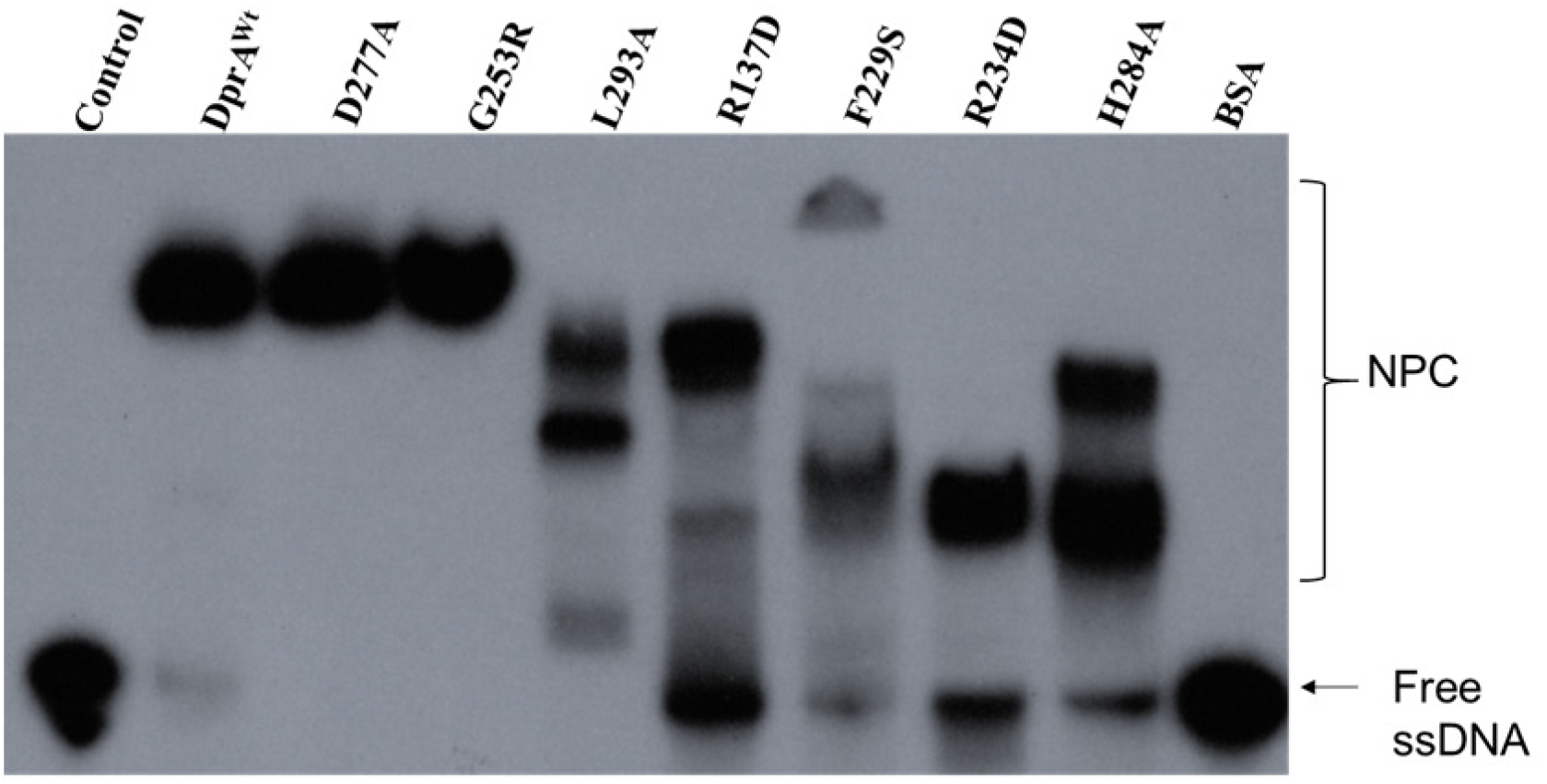
DNA binding activity of DrDprA and its mutant. Recombinant wild-type DrDprA and its different mutants [DprA-RecA interaction (G253R, and D277A), DprA-DprA interaction (L293A, and H284A), and interaction with DNA (R137D, R234D, and F229S] proteins were purified and incubated with [^32^P] labeled 167mer single strand DNA in a reaction buffer as described in methods. Reaction mixtures were separated on 8% native PAGE followed by gels drying and autoradiograms was developed. Experiments repeated three times and were reproducible.

### 5) DrDprA protects DNA from nucleases and limits the SSB inhibitory effect during SER

DprA binds to transforming DNA and protects it from nucleolytic degradation. The stability and half-life of incoming DNA get reduced in *dprA* mutant [12, 48, 49]. Therefore, the DNA protection from nuclease was checked in the presence of a different concentration of purified DrDprA. The DrDprA (0.1 to 1.6 μM) was able to protect the DNA in DrDprA-DNA complex from DNAaseI and T5 Exo degradation (Fig. 7). This property of DrDprA was found to be similar to HpDprA. *In vitro* treatment of HpDprA-ssDNA complex with ExoT, ExoIII, RecJ, T7 Exo, mung bean endonuclease, and DNase I did not show degradation of complexed DNA [42].

**Figure 7.**
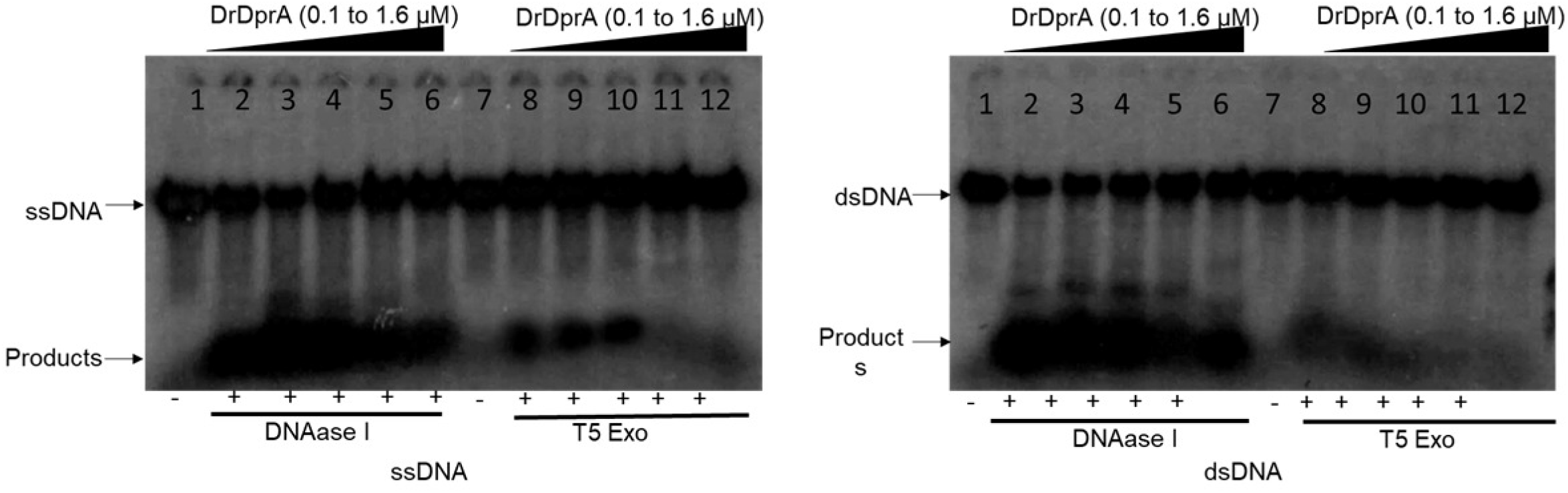
Nuclease protection assay. ^32^P-labelled ssDNA or dsDNA (0.2 nM) either alone or pre-bound with increasing concentrations of DrDprA (0.1, 0.2, 0.4, 0.8 and 1.6 μM, Lane 2 to 6, and Lane 8 to 12) was incubated for 30 min with 1 unit of DNAaseI/ T5 Exo. Lane 1 and 7: DNA alone.

It has been shown that the SsbB and SsbA proteins of *B. subtilis* bind and melt secondary structures in ssDNA but limit RecA nucleation onto DNA. DprA of *B. subtilis* physically interacts with RecA to facilitate the RecA nucleation, and filament extension on SsbB/SsbB-SsbA coated ssDNA by dislodging SSB proteins and thus assists RecA-mediated DNA strand exchange in the presence of both SSB proteins [17, 45]. Here we tested the DrDprA ability to facilitate DNA strand exchange reaction catalyzed by DrRecA in the presence of *E. coli* SSB. The *E. coli* SSB has similar overall protein architecture and ssDNA-binding surface as that of *B. subtilis* SsbB protein [50]. The pre-incubation of ssDNA with SSB (0.3 to 1.2 μM) inhibits the DrRecA catalyzed SER. However, when the SSB-ssDNA complex was incubated with 1 to 4 μM DrDprA followed by addition of RecA and homologous dsDNA, the DNA strand exchange product yield improved substantially (Fig. 5D), indicating a strong possibility of DrDprA counterbalancing the inhibitory effect of SSB in SER reactions. Thus, DrDprA functions nearly similar to the function of BsDprA and RecO protein in the displacement of SsbB/SsbB-SsbA proteins from ssDNA. It has been explained that once the action of DprA/RecO protein gives RecA access to ssDNA, the RecA nucleoprotein filament elongation displaces SSB and enables RecA-mediated DNA strand exchange [17].

## Discussion

Bacteria can take up extracellular DNA (eDNA) by natural transformation (NT) through a well-programmed multistep process coordinated by the multiprotein complex [6, 9]. Many bacteria like *D. radiodurans, H. pylori, and N. gonorrhoeae* have constitutive competence systems and can uptake eDNA throughout their growth phase [26, 27, 41, 42]. DprA (DNA processing protein A) is a transformation-specific recombination mediator protein (RMP) that plays a very significant role in NT in the majority of bacteria [51] and the absence of this gene significantly reduces the transformation efficiency of both chromosomal and plasmid DNA [27, 52–54]. The NT helps bacteria to survive under adverse conditions and contributes to genetic diversity in bacteria. *D. radiodurans,* a radioresistant bacterium, shows natural competence [22, 26, 27] which has been attributed to its genetic diversity and thereby possible role of natural competence in radioresistance cannot be ruled out. *dprA^-^* mutant of *D. radiodurans* showed substantial losses, and the reduction of 160 and 21-fold transformation frequency of chromosomal and plasmid DNA was observed, respectively [27]. The effects of *dprA* deletion on the radioresistance and DSB repair reported earlier [27] conflict with our studies and need more studies to conclude. Therefore, functional characterization of DrDprA would be worth understanding the physiological role of this protein in *D. radiodurans*. Here, we have found evidence to suggest that DrDprA is largely similar to DprA from most other bacteria, except it shows nearly equal affinity to both ssDNA and dsDNA. DprA proteins of many bacteria species have shown a higher affinity towards ssDNA than dsDNA, except HpDprA, which has shown moderate higher affinity for the ssDNA over dsDNA [42]. The structure-function studies of HpDprA and SpDprA identified the potential residues crucial for self-interaction, interaction with DNA, and RecA protein [19, 42]. Full-length HpDprA and its RF domain can dimerize *in vitro,* suggesting that the residues involved in dimerization are located in the N-terminal domain [55]. However, SpDprA forms tail-to-tail dimers and requires the DML1 domain, and this type of dimerization is crucial for nucleoprotein complexing with ssDNA *in vitro* [19]. The sequence alignment studies by Lisboa and colleagues, 2014 identified two motifs in DprA family proteins with essential roles in ssDNA binding [46]. Motif 1 (‘G-S/T/A-R’) is located in the loop (b3–a3) region, and corresponding residues are Gly50–Arg52 in HpDprA. The second motif 2 (‘F/L/Y-X-X-R-N/D’) is located in helix α6 and corresponds to residues Phe140–Asn144 in HpDprA, and these two motifs are considered to be signatures in the DprA domain that binds to ssDNA [56]. MSA analysis of DrDprA with SpDprA, HpDprA, and RpDprA revealed the conserved residues for the DprA-RecA interaction (I287, G253, and D277), DprA-DprA interaction (L293 and H284), and interaction with DNA (R137, R234, and F229). The amino acid residues Arg137 (R137) and Arg232 (R232) of DrDprA lying in DNA binding motif 1 and 2 as identified by Lisboa and colleague, 2014 respectively [46]. Amino acid residues contributing to oligomerization and interaction with DNA and DrRecA were found to be largely conserved across DprAs. DrDprA undergoes oligomerization and interacts with DrRecA. Further, DrDprA supports the DNA strand exchange reaction of DrRecA. The RecA-DprA interaction has been shown for many bacterial species (*S. pneumoniae*, *R. palustris* and *B. subtilis)* but not for *H. pylori* [17, 18, 20, 45, 46]. RecA specifically interacts with DprA, which helps RecA load on SSB-coated ssDNA [52]. DrDprA undergoes oligomerization and interacts with DrRecA *in vivo.* DNA binding activity and oligomerization functions of DrDprA were essential for its support in SER by DrRecA. The perturbation of functional DrDprA and DrRecA interaction with oligomer and DNA binding mutant of DrDprA is not surprising. It has been demonstrated earlier in the case of SpDprA with its cognate RecA [19]. Further, it has been shown that DrDprA dislodge SSB bound to ssDNA and thus removes the inhibitory effect of SSB in SER reaction by RecA [57]. DrDprA bindings with ssDNA and protects from nucleolytic degradation. The DNA protection ability of the DprA protein was also reported for the SpDprA, NmDprA, and BsDprA [20, 41, 45]. Together, our results of biochemical characterization of DrDprA suggested that, like other DprA proteins, DrDprA has functional conservation of all typical DprA properties. Yet, it has some dissimilar properties, like it has all three domains (SMF, RF, and DML1) instead of two domains known for most DprA except NmDprA and RpDprA. On a functional basis, DrDprA has strong dsDNA binding properties along with ssDNA binding. dsDNA binding property may suggest the more diverse function of DrDprA in DNA metabolism, including natural transformation.

## Materials and methods

### 1) Bacterial strains, growth medium, and plasmids

Wild type bacterium *D. radiodurans* R1 (ATCC 13939) and its mutant were *grown in* TGY medium (1% Bacto tryptone, 0.1% glucose, 0.5% yeast extract) with appropriate antibiotic as described earlier [28]. *E. coli* Novablue strain was used for the cloning and maintenance of plasmids. *E. coli* BTH101 (lacking cyaA, referred here as BTH101) was used to determine *in vivo* protein-protein interaction studies by co-expressing the cloned gene on BACTH plasmids and grown at 30°C [29]. Recombinant proteins were purified from overexpressed Top10 *E. coli* cells. pUT18, pKNT25, and pBAD plasmids and their derivatives were maintained in *E. coli* cells (Nova blue) in the presence of the required antibiotics. Various molecular biology techniques and their working protocols were used as described [30]. Antibodies against the T18 (SC-13582) domains of CyaA protein of *Bordetella pertussis* were procured from Santa Cruz Biotechnology, Inc., and the Anti-His antibody was purchased from New England Biolabs, USA. Molecular biology grade enzymes, chemicals, and salts are mainly procured from Sigma Chemicals Company, USA; Roche Biochemicals, Mannheim, Germany; New England Biolabs (USA), and Merck India Pvt. Ltd., India. Radiolabeled [^32^P]-nucleotides supplied by the Board of Radiation and Isotope Technology (BRIT), Department of Atomic Energy, India. Bacterial strains, plasmids, and primers used in the current study are listed in Table S1 (for reviewers’ information only).

### 2) Recombinant plasmids construction and protein purification

Table S1 (for reviewers’ information only) has a list of plasmids and primers used in this study. The transnational fused DrDprA with T18 tag at C-terminus referred to as 18DrDprA encoded by *pUTdprA* plasmid. pRadHisRecA, pUT18recA, and pKNT25recA plasmid were constructed before and transformed into BTH101 *E. coli* cells along with pUT*dprA* plasmid [31]. The expression of fusion proteins in BTH101 cells was confirmed by Western blotting using antibodies specific to the C18 tag and polyhistidine tag. T18 tagged DrDprA cloned in pVHSM shuttle vector at NdeI-XhoI sites for co-immunoprecipitation studies in *D. radiodurans.* Both pVHSVI*dp/A* and pRadHisRecA co-transformed to *D. radiodurans* cells, and coimmunoprecipitation studies were done as mentioned in section 3 of material and methods. Recombinant DrDprA and its various mutants were overexpressed in *E. coli* Top10 cells using 0.2% v/v arabinose. Proteins were purified as described previously [32, 33]. In brief, *E. coli* Top10 cells expressing recombinant proteins were harvested after 3h post-induction by arabinose. The cell pellet was suspended in buffer A (20mM Tris-HCl pH 7.6, 300 mM NaCl, 10% glycerol), 0.5 mg/ml lysozyme, 1mM phenylmethylsulfonyl fluoride (PMSF), 0.03% NP-40, 0.03% Triton X-100, and 10% glycerol and incubated at 37°C for 30 min. Protease inhibitor pellets, NEB make were added to the reaction mixture, and lysis of the cells was done using sonicated for 10 min using 5-second pulses with intermittent cooling for 10 seconds at 35% amplitude. Cell lysate was centrifuged at 12,000 rpm for 30 min at 4°C. NiCl_2_ charged-fast-flow-chelating Sepharose column (GE Healthcare) was equilibrated with buffer A (20mM Tris-HCl pH 7.6, 300mM NaCl, 10% glycerol), and clear cell extract was loaded onto it. The column was washed with 20 column volumes of buffer A containing 20 mM imidazole until detectable proteins stopped coming from the column. Recombinant column-bound protein was eluted with buffer A containing 250mM imidazole. Fractions analyzed on SDS-PAGE, and those containing nearly pure proteins were pooled. Recombinant proteins were further purified on Q-sepharose, MonoQ, and Superdex-200 columns. Protein fractions free from detectable nuclease contamination and having more than 95% purity were pooled and precipitated by ammonium sulphate precipitation, followed by dialysis in buffer B (10mM Tris-HCl pH 7.6, 50mM KCl, 50% glycerol, and 1mM PMSF) and stored at −20°C Proteins.

### 3) Protein-protein interaction studies, Western blotting, co-immunoprecipitation

As detailed elsewhere, *ex vivo* protein-protein interaction studies were done using *a* bacterial two-hybrid system (BACTH) [34, 35]. BTH101 *E. coli* cells were transformed with pUT18recA, pUT18dprA, and pKNT25recA plasmids expressing target proteins with T18 tags or T25 tags at the C-terminus. Empty vectors (pUT and pKNT) in BTH101 cells were used as controls. The cells spotted on LB agar plates containing 5-bromo-4-chloro-3-indolyl-β-d-galactopyranoside (X-Gal) (40μg/ml), IPTG (0.5mM), and antibiotics as required. Plates were incubated at 30°C for 12hr and the appearance of white-blue colored colonies recorded. For the western blotting and co-immunoprecipitation studies, pVHSM*dprA* and pRadHisRecA plasmids expressing C-18 tag (18DrDprA) and His-tag (HisRecA) fusion proteins were cotransformed to *D.radiodurans*. The recombinant cells co-expressing these proteins were induced with 0.5mM IPTG and harvested cells washed with 70% ethanol followed by lysed in the buffer (50mM Tris base, 150mM NaCl, 5mM EDTA) containing 0.5% Triton X-100, 1mM PMSF, 1mM dithiothreitol (DTT), 0.5mg/ml lysozyme, and 50μg of a protease inhibitor cocktail tablet. After sonication, the clear cell-free extracts (CFE) were obtained by centrifugation at 12000Xg for 30 min. CFE used for immunoprecipitation using polyclonal antibodies against the Anti-His tag antibody, and precipitated immunoprecipitates were separated on a 10% SDS-PAGE gel, and protein transferred to polyvinylidene difluoride (PVDF) membrane, and hybridized with monoclonal antibodies specific to T18 tag. Antimouse secondary antibodies conjugated with alkaline phosphatase was used to detect color signal formed by BCIP/NBT (5-bromo-4-chloro-3-indolyl phosphate/nitroblue tetrazolium) substrates (Roche Biochemical, Mannheim, Germany).

Surface Plasmon Resonance (SPR; Autolab Esprit, Netherland) used to study the interaction of DrDprA with DrRecA. 2.5μM DrDprA protein was immobilized on a bare gold sensor chip employing EDC-NHS chemistry (Autolab ESPIRIT SPR User manual) at 20°C. DrRecA (0.5 to 3 μM) was used in the mobile phase (20 mM Tris (pH 7.6) and 1mM MgCl_2_). After deduction of mobile phase buffer controls, the data were processed using the Autolab kinetic evaluation software (V5.4) and plotted after curve smoothening using Graphpad PRISM software.

### 4) DNA strand exchange reaction

Extended homology-dependent RecA-dependent DNA strand exchange was carried out as described earlier [31, 36]. The DrRecA-dependent DNA strand exchange reaction was carried out at 37°C using M13mp18 circular ssDNA and linear dsDNA. Reaction carried out in buffer (25 mM Tris-acetate, 1 mM DTT, 5% glycerol, 3 mM potassium glutamate, 10 mM magnesium acetate, and an ATP-regenerating system (10 units/ml of pyruvate kinase/3.3 mM phosphoenolpyruvate or 10 units/ml creatine kinase/12 mM phosphocreatine). DNA, SSB, ATP, DrRecA, and DrDprA protein concentrations are indicated for each experiment. Reaction initiated with a pre-incubation of ssDNA with DrRecA protein at 37°C for 5 min, followed by the addition of ATP and SSB protein. After incubation of 5-min, linear duplex DNA was added to start the DNA strand exchange reactions. DrDprA protein was added before the addition of dsDNA (wherever required). For the checking inhibitory effect of SSB in SER reaction, SSB (0.3 to 1.2 μM) added before adding DrRecA. The reactions were stopped by adding 5μl of stop solution (0.125% bromophenol blue, 25mM EDTA, 25% glycerol, 5% SDS, and 0.5μg proteinase K), and samples were incubated for another 10 min at 37°C. Samples were electrophoresed in a 0.8% agarose gel with TAE buffer. Gel stained with ethidium bromide and photographed in Gel doc system (Syngene).

For the oligo-based DNA strand exchange reaction, 0.5μM DrRecA incubated with 167mer (2.5μM nucleotides) in 10μl of buffer (25mM Tris-HCl, pH 7.5, 1mm DTT, 2.5mM MgCl_2_, 25mM KCl) containing 1mM ATP for 5 min., after this ^32^P-labeled 40mer dsDNA oligonucleotide (2.5 μM nucleotides) added. DrDprA protein was added as and when required with indicated concentration. To terminate reactions, stop solution (0.125% bromophenol blue, 25mM EDTA, 25% glycerol, 5% SDS, and 0.5μg proteinase K) added and incubated at 37 °C for 10 min. The samples were analyzed on 10% PAGE, the dried gel was exposed to X-ray film, and an autoradiogram was developed.

### 5) Cloning and site-directed mutagenesis

*D. radiodurans* genomic DNA was prepared as described previously. DrDprA coding sequence was PCR amplified using gene-specific primer (Table S1, for reviewers’ information only) and cloned at *XhoI* and *EcoR1* sites in the pBAD vector. The resultant plasmid pBAD*dprA* was used for site-directed mutagenesis of DrDprA. The putative amino acids of DrDprA responsible for the DprA-RecA interaction (G253, and D277), DprA-DprA interaction (L293, and H284), and interaction with DNA (R137, R234, and F229) were selected and site directed mutagenesis (SDM) done as following the kit manufacturers protocols (New England Biolab, USA). The *in vitro* mutagenesis was confirmed by sequencing. The resultant pBAD plasmids expressing G253R, D277A, L293A, H284A, R137D, R234D, and F229S mutants of DrDprA were named as pBADG253R, pBADD277A, pBADL293A, pBADH284A, pBADR137D, pBADR234D, and pBADF229S respectively. These plasmids were transformed into *E. coli* Top10 cells for expression of recombinant proteins.

### 6) DNA binding assay

As described earlier, the DNA binding activity of DrDprA and its mutant derivatives were checked using an electrophoretic gel mobility shift assay (EMSA) [33]. In brief, 40bp or 167bp nucleotide long random sequence oligonucleotide (Table S1, for reviewers’ information only) was used as ssDNA substrate. dsDNA substrate was made by annealing it with its complementary strand (Table S1, for reviewers’ information only). Both ssDNA and dsDNA were labeled with [^32^P] γATP using polynucleotide kinase and purified by the G-25 column. The 2nM labeled probe (ssDNA and dsDNA) was incubated with increasing concentrations of both DrDprA in 10μl of reaction mixture containing 10mM Tris-HCL, pH 7.5, 50mM NaCl, and 1mM DTT for 10 min at 37°C. Products were analyzed on a 10% native polyacrylamide gel, dried, and recorded signals by autoradiography. DNA band intensity in free form or bound to protein was quantified using Image J software. The percent bound fraction of DNA was plotted against protein concentration using Graphpad Prism 5. The Kd for curve fitting of the individual plot was determined by the software working on the principle of least squares method applying the formula Y= Bmax *[X]/Kd+[X], where [X] is the protein concentration and Y is the bound fraction. The DNA fraction bound to protein was plotted as a function of protein concentration using Graphpad Prism 5. to know the DrDprA DNA binding preference between ssDNA and dsDNA the competition assay was performed where the binding of 0.2 nM of ssDNA/dsDNA was challenged with either unlabelled homologous ssDNA and dsDNA (10nM to 80nM) respectively. LogKi was calculated by curve fitting using nonlinear regression of competition binding equation of one site Fit Ki in Graphpad PRISM software.

